# Genetic diversity by ISSR of two endemic quillworts (*Isoetes* L.) species from Amazon Iron Rocky Outcrops, *Isoetes cangae* e *I. serracarajensis*

**DOI:** 10.1101/635771

**Authors:** Mirella Pupo Santos, João Victor da Silva Rabelo Araújo, Arthur Vinícius de Sant’anna Lopes, Julio Cesar Fiorio Vettorazzi, Rodrigo Nunes da Fonseca, Emiliano Nicolas Calderón, Marcela Santana Bastos Boechat, Fernanda Abreu Santana Arêdes, Messias Gonzaga Pereira, Naiara Viana Campos, Fernando Marino Gomes dos Santos, Taís Nogueira Fernandes, Rodrigo Lemes Martins, Francisco de Assis Esteves

## Abstract

Two lycophytes endemic species have been recently described at the State of Pará, in the Amazon forest located in the North of Brazil. Genetic diversity and population structure of *Isoetes cangae* and *I. serracarajensis* were investigated through ISSR molecular markers. These analyses aim to establish strategies for future attempts for species conservation. From sixteen primers, 115 gel bands were identified from which 87% were polymorphic. A high level of polymorphic loci (81,74 % e 68,48 %) and a high Shannon index for intra populational genetic diversity was observed for each species (Sh=0.376 e 0.289) *I. cangae and I. serracarajensis*, respectively. The largest genetic diversity of both species relies in their own populations. The coefficient of genetic differentiation between population areas (G_ST_) was higher in *I. serracarajensis* (0.5440) than in *I. cangae* (0.2250). Gene flow was high between *I. cangae* populations (1.7142) and very low in *I. serracarajensis* (0.4190). Principal Component Analysis (PCoA) showed that individual plants were allocated into species-specific and population groups. Overall, the results further show that *I. serracarajensis* (0.5440) and *I. cangae* are two species with considerable genetic variation. These results should be considered for effective conservation strategies of both species.

## Introduction

The genus *Isoetes* L. is characterized by approximately 200 species of vascular plants with molecular dating studies tracing its evolutionary roots to the Devonian, in the Paleozoic^1^. This genus has survived to three mass extinctions probably due to adaptation of large environmental changes over hundreds of million years. *Isoetes* are considered small-sized heterosporate lycophytes ^2^ ^3^, which can display a great diversity of metabolic strategies ^4^. These lycophytes can be associated to humid environments such as seasonally flooded plains and oligotrophic lakes.

Low nutrient availability is a common feature of *Isoetes* habitat. This adaptation is correlated to the activation of the CAM metabolism, which guarantees that the photosynthetic efficiency is high even in regions of low carbon concentration^5^. Some aquatic species from the genus, for example *I. lacustres* induce oxidation of soluble nitrogen compounds such as nitrate by the release of oxygen from the roots increasing the process of denitrification and the local redox potential^6^ ^7^ ^8^. *Isoetes* species might also act as oligotrophic agents promoting phosphorus immobilization^9^ ^10^.

*Isoetes* genus also shows a variety composition of ploidy, which has been suggested to be an important and determinant feature for adaptation in several region of the planet, thus the genus can be considered almost as cosmopolitan^11^ ^12^. Sexual and asexual reproduction by self-fertilization and cross-fertilization has also been reported, besides apomixis^13^ ^14^ ^15^.

Apomixis reproduction has been described when megaspores germinate asexually in the mother plant. This phenomenon occurs in populations which have lost the capacity of sexual reproduction after population bottlenecks or chromosomal imbalance ^15^ ^16^. In these species the lifetime of megaspore and, consequently, the rate of asexual reproduction can be correlated with the ploidy level and with the imbalance number of ploidy ^17^ ^18^. The sexual reproduction by self-fertilization is limited by non-synchronous maturation of microspores and megaspores and which is more common in polyploidy populations due to the effect of additional genome buffering, reducing the effect of depression by endogamy ^19^.

Sexual reproduction by cross-fertilization results in larger gene flow and is related to the geographic distance from the genitors and the presence of spore dispersing agents ^20^ ^21^ ^22^ ^14^ ^23^. The mating system directly influences the genetic variation of each species; thus it is a fundamental factor for population establishment and survival. Conservation studies of *Isoetes* genus and other plant groups requires proper knowledge of the mating system^24^ ^25^ ^26^. Since *Isoetes* species can survive as asexual as well as sexual species,^27^ ^18^ the specific genetic diversity generated is associated to the inherent properties of population colonization and isolation. This diversity can be accessed by molecular markers such as the *Inter Simple Sequence Repeat* (ISSR), a dominant marker of multi allelic nature, high reproductibility and large genome coverage. ISSR reveals a large number of polymorphic fragments and, thus, constitute a suitable marker of intra and inter populational genetic diversity for *Isoetes* ^24^ ^25^.

The species *Isoetes cangae* and *I. serracarajensis*^2^ targets of the current study were recently described from Amazon ferruginous fields of Serra dos Carajás (Figure 1), southeast of the state of Pará, northern Brazil. The species *I. cangae* is characterized as an aquatic plant with endemism restricted to a lagoon of the ferruginous plains of the southern region of the Serra dos Carajás. *I. cangae* restricted endemism and aquatic condition contrasts with its congener *I. serracarajensis* that is found in several ferruginous plateaus of southeastern Pará, associated with seasonally flooded environments. Recent cytogenetic studies revealed that *I. serracarajensis* is tetraploid and *I. cangae* is diploid^28^. *I. serracarajensis* displays CAM and C3 metabolism (unpublished data), maybe an important adaptation to its amphibian lifestyle, while *I. cangae* shows CAM metabolism (unpublished data) being restricted to oligotrophic aquatic environments. The species *I. cangae* presents sexual reproduction, demonstrated by *in vitro* fertilization and by self-fertilization (unpublished data), while *I. serracarajensis* reproduction so far has been unable to establish *in* vitro methods or to demonstrate its reproduction type.

**Figure 1.**
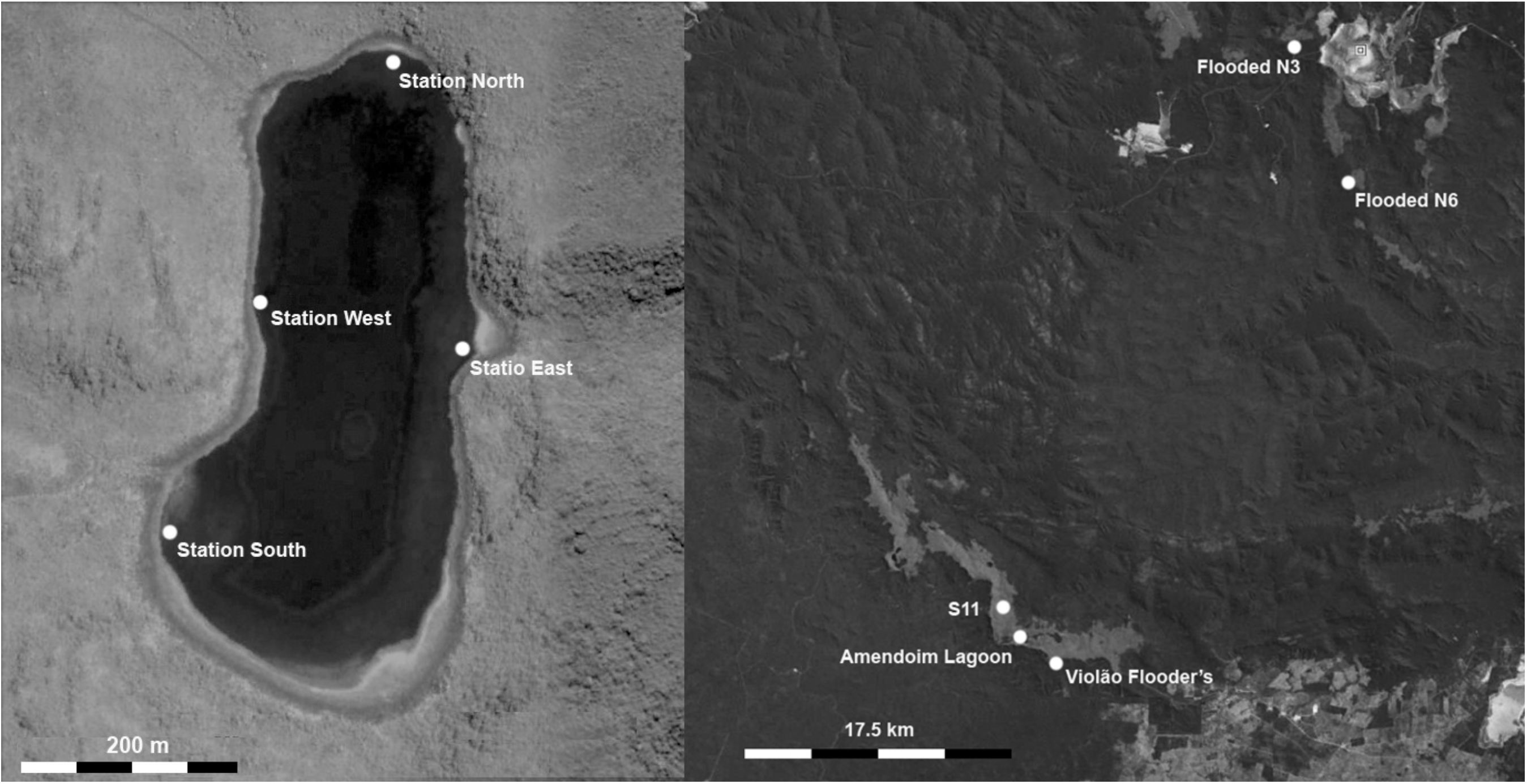
Iron Rocky Outcrops from Serra dos Carajás, Pará State, Brazil. (A) Four collection points of *Isoetes cangae.* (B) *I. serracarajensis* was gathered at different regions in the Serra dos Carajás, two locations on the ferruginous plateaus north of the Serra dos Carajás, flooded N3 and N6, and two locations on the ferruginous plateaus of the southern portion, called S11D marsh and Tapir’s Bath.

All this scenario of metabolic and lifestyle differences among both species provides a very interesting comparative picture, which could be reflected in the genetic diversity of populations. Thus, the current study aimed to estimate the genetic and structural variation of the population of *I. cangae* and *I. serracarajensis.* This information will enable the establishment of conservation strategies of the genetic resources of these species that guarantee their success and perpetuation in new environments.

## Results

### ISSR polymorphism

Sixteen primers produced a total of 115 loci. One hundred one (101) were polymorphic loci (mean of seven loci per primer) which correspond to almost 87% of the whole sample. The most informative primer was LB6 with nine polymorphic loci (Table 2). The amplified fragments ranged from 100 to 2000 bp. *Isoetes cangae* species showed 81.74% of polymorphic loci and a high diversity index (He = 0.245) (Table 3). The ICO area displayed the highest percentage of polymorphic locus (64.35%) by primers (PLP), while the lowest percentage was obtained with the ICS, 33.04%. The expected heterozygosity (He) values ranged from 0.187 in ICO to 0.096 in ICS. The ICO and ICN area presented higher number of rare loci (Table 3). The populations of *I. serracarajensis* showed 63.48% of polymorphic locus and diversity index He of 0.187. The population with the highest PLP was ISV (52%) and the lowest was the IS11 (18%). Among the populations, ISV displayed the highest expected heterozygosity value (0.164), while IS11 displayed the lowest (0.043). The ISN3 and IS11 also showed five rare loci, while ISN6 and ISV had three (Table 3).

### Genetic structure of population

Population structure of both species was evaluated by a series of parameters. The species *I. cangae* presented a high diversity index at the species level, Sh = 0.376, G_ST_ equal to 0.2250 and high gene flow (1.7142). The populations of *I. serracarajensis* displayed high to medium values of diversity index of Shannon (Sh=0.289), G_ST_ equal to 0.5440 and low genetic flow (0.4190) (Table 3).

From the dissimilarity matrix, based on the genetic distances of Nei (1968), a dendogram was generated to investigate population clustering (Figure 2). The results show the separation in two well-defined clusters between *I cangae* species (ICN areas, ICS, ICL, ICO) and the populations of the species *I. serracarajensis* (ISN3, ISN6, ISV, IS11). Comparison of *I. cangae* gathering sites shows that the ICO forms a separate group from the other populations of the same species, possibly due to the number of polymorphic and rare loci (Figure 2, Table 3) of the specimens sampled compared to other areas.

**Figure 2.**
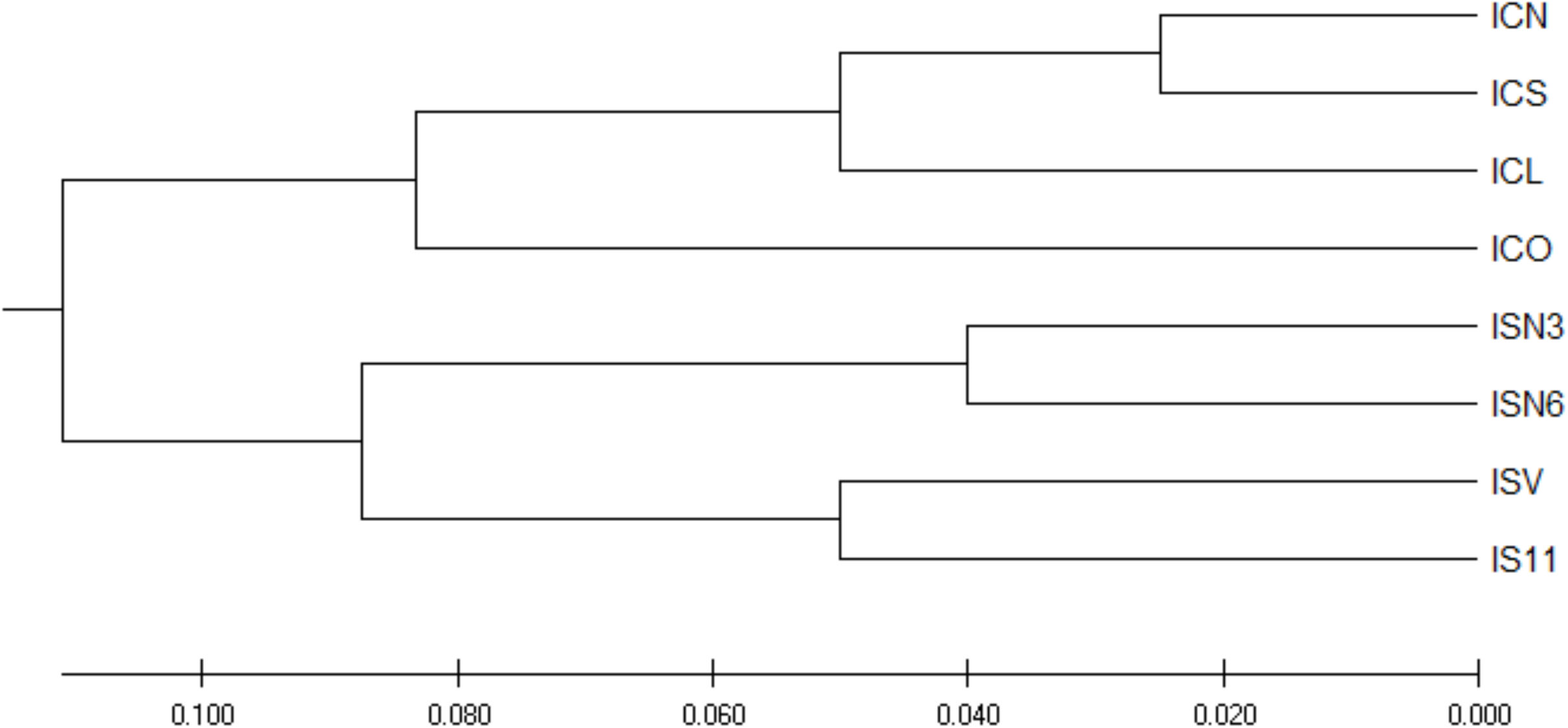
Dendrogram obtained from the dissimilarity matrix based on the genetic distances of Nei. ISSR fragments detected in four populations of *I. cangae* and in four populations of *I. serracarajensis*.

A matrix of genetic distance and geographic distance was constructed (Table 4). Geographic distances showed no correlation with the genetic distance of Nei for the species *I. cangae* (Figure 2A). The values of genetic distance of Nei among the populations of *I. serracarajensis* generated clear groups of populations. ISN3 and ISN6 populations which occur in the same region (Serra Norte) were located in the same group, while ISV and IS11 populations in another group (Serra Sul), however, statistical analysis did not show a correlation between the geographic area of sample collection and the genetic distance values of Nei (Figure 3B).

**Figure 3.**
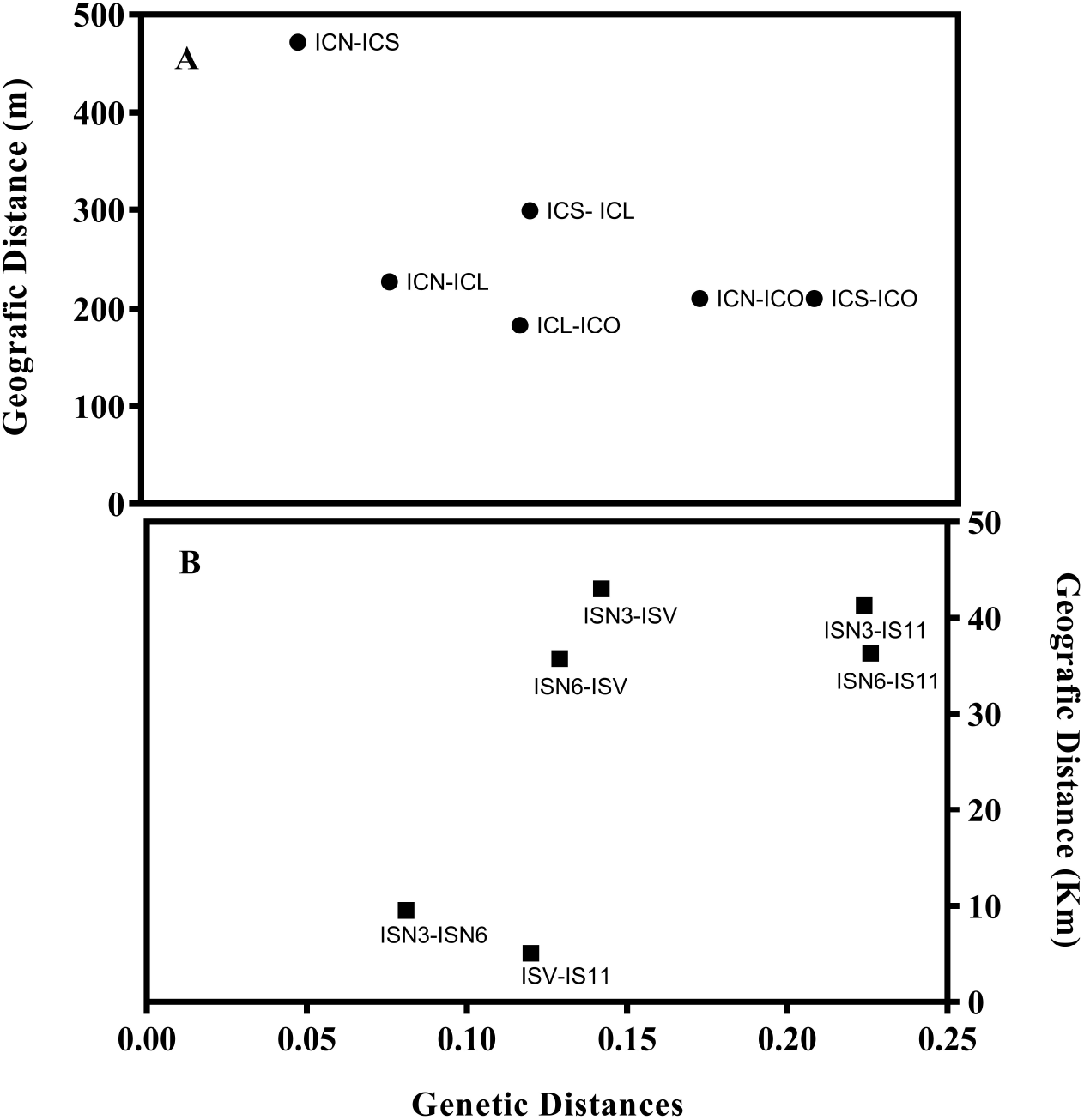
Relationship between pairwise geographic and pairwise genetic distance. Graphic among the four local populations of *I. cangae* (A) and *I. serracarajensis* (B). There was no correlation between geographical distance (m) and genetic distance among populations with the Mantel test to populations of *I. cangae* and of *I. serracarajensis* (r= −0.5218 P=0,2972)

### Analysis of the main coordinates and AMOVA

From the analysis of the main coordinates (Figure 4), graphic dispersion of the individuals and allocate them into different groups. This analysis aimed to verify the genetic diversity of the individuals under study. Four large groups were observed; two corresponding mainly to *I. cangae* and the other two groups contains *I. serracarajensis* individuals. The first group includes individuals from the ICN, ICS, ICL and ICO areas; the second group is made of IS11 and *Isoetes cangae* individuals. The third group comprises the populations of ISN6, ISV individuals. The fourth group contains ISN3 and ISN6 specimens. Groups were well defined in relation to the species to which they belong, with the noticeable exception for sample 48 that belongs to the species *I cangae* and was grouped together with group two, constituted by *I. serracarajensis*.

**Figure 4.**
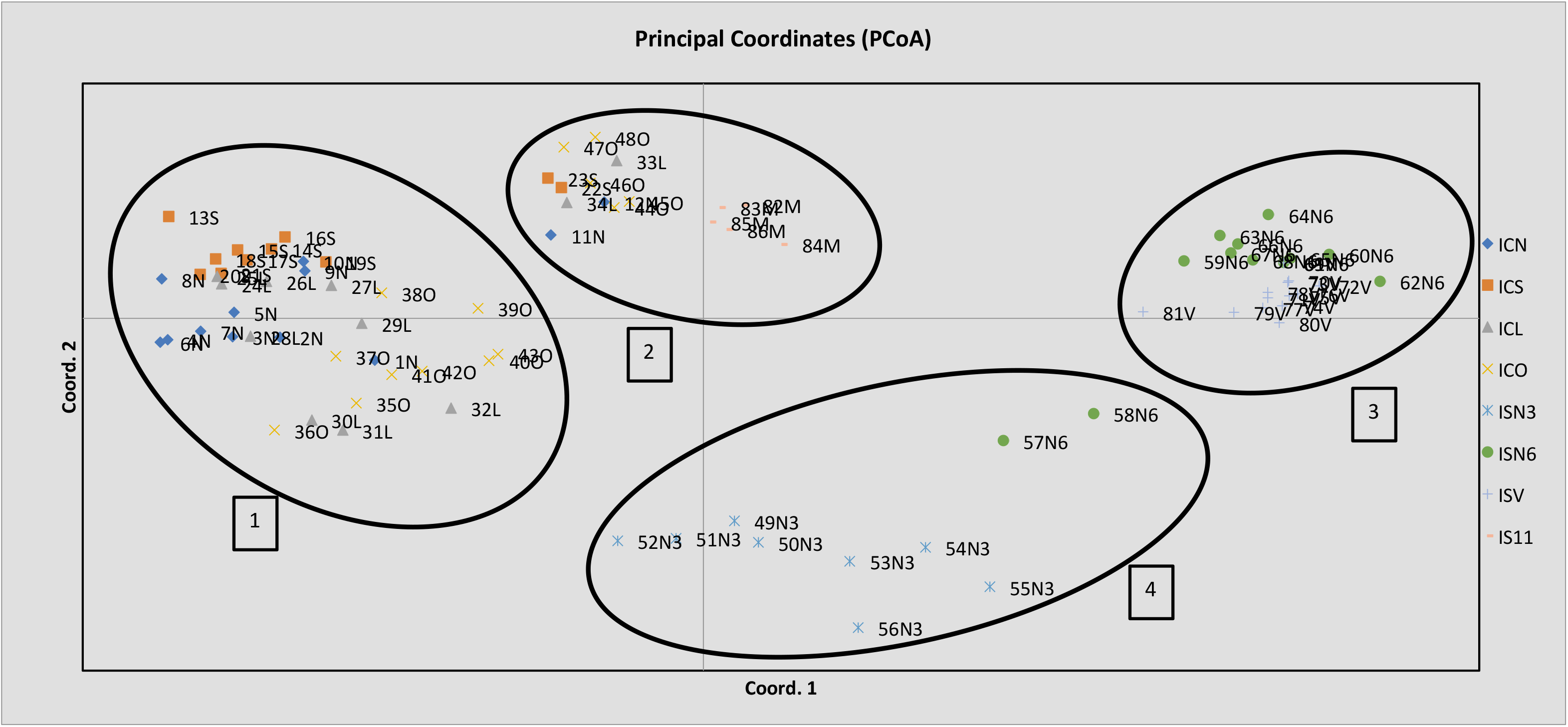
Analysis of the main components of the *I. cangae* and *I. serracarajensis* species. Clusters are delimited by the circles: 1) includes individuals from the ICN, ICS, ICL and ICO areas; 2) IS11 and *Isoetes cangae* individuals; 3) Comprises the populations of ISN6 and ISV individuals; 4) Contains ISN3 and ISN6 specimens.

Quantification of variability was obtained through the analysis of molecular variance (AMOVA). Comparison of total genetic variance accessed by ISSR markers showed that for *I. cangae* 84% was observed within each population and 16% among populations. Genetic variance of *I. serracarajensis* species was 55% within each population and 45% among the populations (Table 5).

## Discussion

The genetic diversity of species is a result of evolutionary processes, including recent changes such as habitat reduction, which may lead to a reduction in genetic diversity, decreasing the surviving potential for a given species. Our study shows high genetic diversity within and between the populations in both species *I. cangae* and *I. serracarajensis*. The analyzes of molecular variance (AMOVA) showed greater diversity within the populations, *I. cangae* (84%) and *I. serracarajensis* (55%) than between population. The greatest diversity within populations can be explained by several factors such as genetic drift, selection and mutation in groups of plants physically isolated^13^ ^29^.

The dispersion of the ciliary microspores of the genus, which reach a few centimeters of the mother plant, favors reproduction between individuals closely located, resulting in higher genetic similarity between neighboring individuals of the same population ^29^. Results of the ISSR markers revealed high rates of polymorphic locus, 81.74% and 68.48%, and high intra-populational genetic diversity (Sh), 0.376 and 0.289, for *I. cangae* and *I. serracarajensis*, respectively. The colonization of individuals, interspersed with areas with low population density, is common in the study area, both for *I. cangae* and *I. serracarajensis*. Thus, the physical distance between these clusters of plants contributes to diversity among them, which may favor the occurrence of genetically differentiated sub-populations ^30^ ^31^. Chen et al. (2005), Kang et al. (2005) and Kim et al. (2005) found similar results suggesting that factors such as genetic drift may influence intra-populational diversity variation, especially in isolated subpopulations.

Greater intrapopulation diversity for *I. cangae* compared to *I. serracarajensis* was observed. The explanation of this observation might be related to the type of reproduction. The sexual reproduction of *I. cangae* observed in the laboratory by our group. Thus far, the reproduction of *I. serracarajensis* remains unknown, due to the difficulty of sexual and asexual reproduction *in vitro*, as well as to the low viability of gametes of the species outside the aquatic environment (unpublished observations). Another possibility relies on different environments in which both species are found. *I. cangae* is an exclusively aquatic plant, and its spores might be transported all over the year. In contrast, *I. serracarajensis* is observed in seasonally flooded terrestrial habitat, thus spore dispersion is limited to the presence of water during rainy periods. Another possibility to explain this difference in genetic diversity is the level of ploidy of the species and the presence of long-lived individuals and overlapping generations (unpublished observations)^13^.

The diversity index among populations (G_ST_) was high in both species*, I. cangae* (0.2250) and *I. serracarajensis* (0.5440). These data are consistent with the results of gene flow between populations, *I. cangae* (1.7142) and *I. serracarajensis* (0.4190). Gentili et al. (2010) observed, through ISSR and AFLP markers, that population of *I. malinverniana* living along a river showed from average to high genetic diversity due to a substantial amount of gene flow among the populations analyzed.

Despite high gene flow and reduced geographic distance between the populations of *I. cangae*, no correlation was observed between the genetic distance indices of Nei (Fig. 2) and the geographic distance for this species (Fig 3A). This may be related to the limited size of the lagoon and the dispersion of the spores by extrinsic factors of high vagility such as fish and other animals (alligators and turtles) recorded for the study area. Other authors demonstrate that the dispersion of spores over long distances may occur between distant groups, but this event is dependent on external factors such as water flow and animals^20^ ^14^ ^23^. The genetic distance of Nei is also not correlated with the geographic distance for the species *I. serracarajensis* (Fig. 3B).

The theory of population genetics predicts correlations between population density and levels of genetic variability so that large populations maintain higher levels of variability than small populations, and that the greater the isolation between populations, the greater the diversity between them^32^ ^33^ ^34^. For *I. cangae* the population ICS, presented higher population density however, revealed lower rates of diversity. These diversity results might be attributed to a reduced distance between individuals, which facilitates spore dispersal and cross-fertilization among genetically similar individuals. In contrast, the ICO population, which presents the lowest population density, showed highest genetic diversity (Table 4), suggest that this area might have been colonized by non-related plants or plants with different genetic backgrounds. Similar results have been published showing that plant density can be inversely correlated to genetic diversity values, since the isolation of individuals can generate fixation and allele loss over time, resulting in high intrapopulation diversification^24^

For the conservation strategies of the species, the genetic structure of the populations and the level of genetic diversity provides a framework for a better understanding of the type of reproduction, guiding the strategies of species preservation. If gene flow between individuals and populations is low, as we observed for *I. serracarajensis*, genetic differentiation within a population might be caused by genetic drift. In this situation, the cross-fertilization between different populations through *in vitro* fertilization, for instance, may increase genetic diversity. ^35^

If gene flow were high among populations, as our data suggests for *I. cangae*, which display higher genetic diversity within the population, it would be interesting to ensure that the greatest diversity is represented if this endemic species is reintroduced into a new habitat. Moreover, our results provide important biological information for future functional studies to unveil the biochemical adaptations behind the endemic quillworts (*Isoetes* L.) species from Amazon ferruginous fields to their specific environments. The knowledge of the current genetic structure of the two species of *Isoetes* allows inferences about past processes and represents an important prerequisite for the prediction of evolutionary responses and effective conservation strategies of the two species.

## Material and Methods

### Plant Material

Specimens from *Isoetes cangae* and *I. serracarajensis* were collected in February 2018 in the Iron Rocky Outcrops from Serra dos Carajás, Pará State, Brazil. *I. cangae* was collected at four points from a lagoon of 12,98 ha arranged on a ferruginous plateau south of the Serra dos Carajás, known as Serra Sul. This is the original location of species description and, thus far, the only known population described. Forty-eight individuals were collected between the northern, southern, eastern and western portions of the lagoon (Fig 1A). *I. serracarajensis* was gathered at different regions in the Serra dos Carajás, two locations on the ferruginous plateaus north of the Serra dos Carajás, flooded N3 and N6, and two locations on the ferruginous plateaus of the southern portion, called Violão flooded and S11 (Fig 1B). The four sampled populations were defined as ISN3, ISN6, ISV and IS11, and a total of 39 specimens were collected. Location information, habitat, area occupied by population (m^2^), categorized population density and number of samples are presented in Table 1.

**Table 1:** Identification of the populations gathered in the state of Pará, northern region of Brazil. POP = Population; LOC: Location; CD: Population Code; Latitude/Longitude (Lat / Log); Habitats (Habt): Aquatic (Aqu) and Aqu/Terrestrial (Ter); AOPE: Area Occupied by Population (m^2^); DENS: Population density - categorized (each specimens/m2); Sample Size: Number of samples.

| POP | LOC | CD | Lat/Lon | Habt | AOP<br>E | DENS | Sample<br>Size |
| --- | --- | --- | --- | --- | --- | --- | --- |
| *I. cangae* north | Serra<br>Sul | ICN | 6°23'47,95"/50°22'17,05" | Aqu | 1.536 | + | 12 |
| *I. cangae* south | Serra<br>Sul | ICS | 6°24'03,73"/50°22'23,27" | Aqu | 1.041 | +++++ | 11 |
| *I. cangae* east | Serra<br>Sul | ICL | 6°23'57,06"/50°22'14,37" | Aqu | 1.188 | ++++ | 11 |
| *I. cangae* west | Serra<br>Sul | ICO | 6°23'55,84"/50°22'21,01" | Aqu | 782 | + | 14 |
| *I. serracarajensis* N3 | Serra<br>Norte | ISN3 | 6°02'44,90"/50°12'34,68" | Aq/Ter | 4.340 | + | 8 |
| *I. serracarajensis* N6 | Serra<br>Norte | ISN6 | 6°07'33,97"/50°10'39,43" | Aq/Ter | 99 | ++ | 12 |
| *I. serracarajensis* ISV | Serra<br>Sul | ISV | 6°24'31,39"/50°21'05,38" | Aq/Ter | 4.959 | +++ | 13 |
| *I. serracarajensis* IS11 | Serra<br>Sul | IS11 | 6°22'33,90"/50°23'00,34" | Aq/Ter | 1.561 | ++ | 6 |

#### Estimates of specimen density and distance between collection sites

The density estimate of the *I. cangae* specimens were obtained to understand the processes related to the maintenance of the diversity of this species of restricted occurrence. For density estimation, coverage and number of individuals were performed by free diving at approximately 12 points randomly distributed in each of the four sampling areas. At each point a square of plastic (PVC) of 1 x 1 m, subdivided into four quadrants of 0.25 m^2^, was arranged randomly on a substrate. In each square the coverage of *I. cangae* was estimated and from fifty squares sampled, counting was performed in fifteen. The relationship between specimen density and coverage occupied in these squares was used to calculate the mean density of specimens in all squares sampled.

For *I. serracarajensis* the density estimation was performed by counting the individuals of the sampled spots and an approximate estimate of the collection area to establish a density category of the specimens. Each point sampled had the geographical coordinates (latitude and longitude) recorded in GPS. The measurement of the area occupied by the sampled specimens (AOPEA), and respective distances between sampled areas, was performed in the Google Earth Pro program (Google LLC 2019). To evaluate area size a polygon comprising all the samples of each area was drawn and is represented in m^2^. The smallest distance between the polygons of the different sampling areas was used to establish the distance among them.

### Total DNA extraction and ISSR PCR amplification

Total genomic DNA was isolated from 1 gram of dry leaves. Leaves were macerated using liquid nitrogen (N_2_) and the extraction was performed with the help of the DNeasy Plant Maxi Kit (Qiagen). The DNA was analyze quantify in Nanodrop and 1μg/μl was used for each PCR reaction. Annealing temperatures varied between 48°C and 52°C according to the primer used. Among the twenty tested primers, sixteen were selected for analysis (Table 2). Quantification of genomic DNA was performed by 2% agarose gel electrophoresis.

**Table 2:** Description of ISSR polymorphic primers validated for the genus *Isoetes*. ID: Identification of the primers; Sequences (5’-3 ”); Ta (ºC): annealing temperature; NLP: Number of polymorphic loci; NTL: Total number of loci; PLP%: Percentage of polymorphic loci per primer.

| ID | Sequences (5'-3') | Ring temp. (°C) | NLP | NTL | PLP (%) |
| --- | --- | --- | --- | --- | --- |
| LB1 | (GA)<sub>6</sub>C | 50 | 5 | 8 | 62,5 |
| LB2 | (GT)<sub>6</sub> C | 48 | 4 | 5 | 80 |
| LB5 | (AG)<sub>8</sub>YT | 52 | 7 | 8 | 87.5 |
| LB6 | (AC)<sub>8</sub>(CC)G | 52 | 9 | 9 | 100 |
| LB11 | (GGAT)<sub>3</sub>GA | 50 | 7 | 8 | 87.5 |
| LB14 | (GA)<sub>8</sub> C | 48 | 6 | 7 | 85,7 |
| LB18 | (AGA)<sub>8</sub>YC | 50 | 9 | 10 | 90 |
| LB19 | (AG)<sub>8</sub>YA | 50 | 6 | 7 | 85,7 |
| LB23 | (AC)<sub>8</sub>YG | 48 | 4 | 5 | 80 |
| LB40 | (AC)<sub>8</sub>(CT)T | 48 | 8 | 8 | 100 |
| LB41 | (TC)<sub>8</sub>(AG)G | 52 | 7 | 7 | 100 |
| LB42 | (TC)<sub>8</sub>(AG)A | 48 | 4 | 5 | 80 |
| LB43 | (GT)<sub>8</sub>(CT)C | 52 | 7 | 8 | 87.5 |
| LB48 | (GA)<sub>8</sub>T | 48 | 5 | 6 | 87.5 |
| LB52 | (ATC)<sub>6</sub>C | 48 | 8 | 8 | 100 |
| LB57 | (GA)<sub>8</sub>AC | 48 | 4 | 5 | 80 |
| **Total** |  |  | 100 | 115 |  |

**Table 3:**
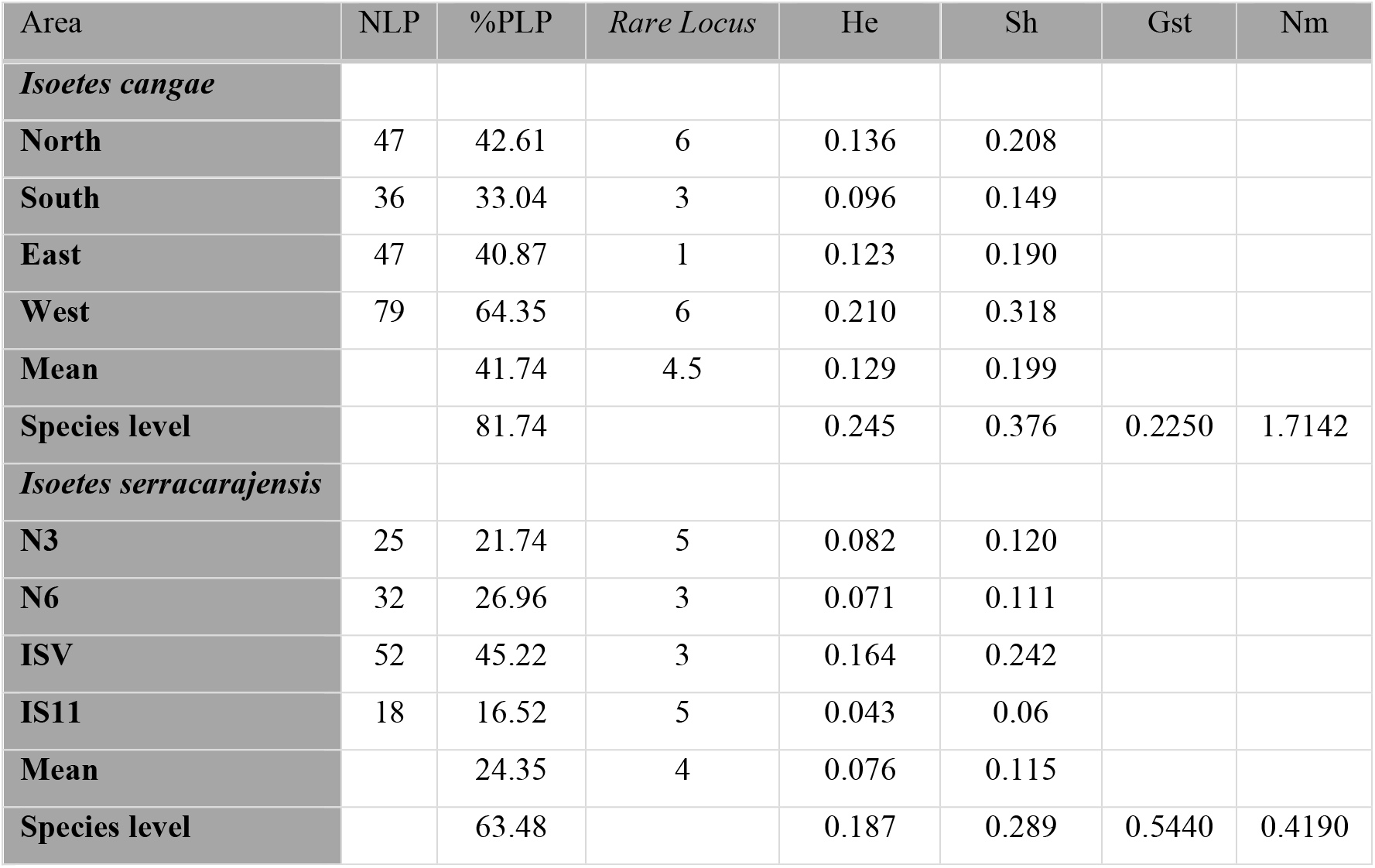
Genetic diversity among the populations of *Isoetes cangae* and *I. serracarajensis*. NLP: Total polymorphic locus; PLP: Percentage of polymorphic loci; Rare Locus; HE: corrected expected heterozygosity of Nei (1978) (assuming the Hardy-Weinberg equilibrium); Sh: Shannon & Weaver Diversity Index. Gst: Proportion of genetic diversity among populations; Nm: Gene flow.

**Table 4:** Genetic distance matrix based on ISSR bands: using pairwise estimated values of Nei genetic distance (below diagonal) and geographical distances (above diagonal) between populations of *I. cangae* and *I. serracarajensis*

| Population | ICN | ICS | ICL | ICO |
| --- | --- | --- | --- | --- |
| ICN | - | 472 | 227 | 210 |
| ICS | 0.953 | - | 299 | 212 |
| ICL | 0.927 | 0.888 | - | 183 |
| ICO | 0.843 | 0.814 | 0.817 | - |
| Population | ISN3 | ISN6 | ISV | IS11 |
| ISN3 | - | 9.546 | 43.003 | 41,250 |
| ISN6 | 0,081 | - | 35,783 | 36.358 |
| ISV | 0,142 | 0,129 | - | 5,020 |
| IS11 | 0,224 | 0,212 | 0,097 | - |

**Table 5.** Analysis of molecular variance of populations *I. cangae* and *I. serracarajensis* via ISSR markers. FV = Sources of Variation; GL = degrees of freedom; QM = mean square; CV = Components of Variance.

| *I. cangae* |  |  |  |  |  |
| --- | --- | --- | --- | --- | --- |
| FV | GL | QM | CV | % | Probability |
| Among pops | 3 | 34.543 | 2,028 | 16 | <0.001 |
| Within pops | 44 | 10,295 | 10,295 | 84 | <0.001 |
| Total | 47 |  | 12.323 | 100 |  |
| *I. serracarajensis* |  |  |  |  |  |
| FV | GL | QM | CV | Percentage | Probability |
| Among pops | 3 | 77.612 | 7.477 | 45 | <0.001 |
| Within pops | 34 | 9.270 | 9.270 | 55 | <0.001 |

### Data Analyses

A binary matrix was constructed from the gel analysis. Presence of bands is indicated by number 1 and absence by number 0. Genetic dissimilarity through the Weighted Index was developed through this binary matrix.

Genotype dispersion by Principal Coordinate Analysis (PCoA), allele frequency analysis, molecular variance analysis (AMOVA), diversity indexes such as Shannon index (I), expected heterozygosity (HE), number of alleles (Na), and effective number of alleles (Ne) were calculated with the help of the software Genalex 6.3. ^36^

A matrix of genetic distance and geographic distance was constructed using GraphPad Prism 7.00^39^. Analyses was performed with the Mantel test to populations of *I. cangae* and of *I. serracarajensis.* Dendogram, one by the UPGMA (unweighted pair group method with arithmetic mean) hierarchical method dissimilarity matrix based on the distance of Nei for the species *I. cangae* and *I. serracarajensis* were developed with the help of the software MEGA (Molecular Evolutionary Genetics Analysis)^37^

Gene flow was calculated using the POP gene version 1.32 program^38^. The correlation between the genetic distance (Nei genetic dissimilarity index) and the geographic distance between the populations of *I. cangae* and *I. serracarajensis* were used to generate a matrix and the correlation between these variables was investigated by the Mantel test using the program Genalex 6.3.

